# Land Use and Endemic Avian Biodiversity on Nusa Penida

**DOI:** 10.1101/2021.06.20.449190

**Authors:** Ashlee A. Abrantes, William D. Brown

## Abstract

Understanding anthropogenic alterations to land use and their effects can inform conservation efforts in tropical biodiversity hotspots. In 2004 the Indonesian Palau Penida Archipelago, off the coast of Bali, was established as an unofficial bird reserve; however, studies of the islands’ land use and avian biodiversity were never conducted and have not been monitored. I surveyed birds across 32 transects in land use categories designated: agriculture, deforested, developed, and forest. Forest transects presented the greatest endemic species richness, but overall, Shannon diversity different significantly among land use categories, particularly forested and deforested. ANOVA indicated exotic bird density was significantly higher than endemic bird density across all transects. Birds serve as a common biodiversity barometer and this study can serve to inform land use management decisions on the Archipelago and throughout reserves and protected areas throughout the tropics.

**Ringkasan:** Memahami perubahan antropogenik untuk penggunaan lahan dan dampaknya dapat menginformasikan upaya konservasi di hotspot keanekaragaman hayati tropis. Pada tahun 2004 Kepulauan Palau Penida Indonesia, di lepas pantai Bali, didirikan sebagai cagar burung tidak resmi; Namun, studi tentang penggunaan lahan pulau-pulau dan keanekaragaman hayati burung tidak pernah dilakukan dan belum dipantau. Saya mensurvei burung di 32 transek dalam kategori penggunaan lahan yang ditunjuk: pertanian, gundul, dikembangkan, dan hutan. Transek hutan menunjukkan kekayaan spesies endemik terbesar, tetapi secara keseluruhan, keanekaragaman Shannon berbeda secara signifikan di antara kategori penggunaan lahan, terutama yang berhutan dan digunduli. ANOVA menunjukkan kepadatan burung eksotis secara signifikan lebih tinggi daripada kepadatan burung endemik di semua transek. Burung berfungsi sebagai barometer keanekaragaman hayati yang umum dan studi ini dapat berfungsi untuk menginformasikan keputusan pengelolaan penggunaan lahan di Kepulauan Indonesia dan di seluruh cagar dan kawasan lindung di seluruh daerah tropis.

## INTRODUCTION

Biodiversity maintenance is essential to tropical ecosystem structure and function (Cardinale et al. 2012, Allan et al. 2015). With greater than two thirds of all known species existing between the Tropics (Pimm and Raven 2000, Brown 2014), anthropogenic changes in land use, such as agriculture, can negatively affect tropical biodiversity, potentially impairing ecosystem function (Gardner et al. 2009, Moura et al. 2013, Newbold et al. 2015). Natural reserves and other protected areas are conservation tools used to mitigate the potential detrimental effects of land use alterations on biodiversity (Juffe-Bignoli et al. 2014). “Ancillary” reserves are conservation areas established outside of International Union for Conservation of Nature (IUCN) standards or government regulation and are also intended as instruments of biodiversity conservation (Borrini-Feyerabend et al. 2013), such as the one established in 2004 on Indonesia’s Palau Penida Archipelago. This study examined avian presence and quantities across distinct anthropogenic land uses on the Palau Penida Archipelago. Community structure assessing unique species, habitat parameters, and endemic status were taken into account. Patterns illuminated diminished endemic populations in the fragmented forests of the Palau Penida Archipelago as well as exotic species dominating the landscape.

Nusa Penida, Nusa Lembongan, and Nusa Ceningan are a trio of islands identified as the Indonesia’s Palau Penida Archipelago. The Palau Penida Archipelago’s juxtaposition of proximity and isolation from mainland Bali created a complex history of human occupation and land use. By the turn of the 20th century, agriculture dominated the small Indonesian islands. A 1924 Dutch survey of Nusa Penida documented 17,800 ha or 86% of the island had been deforested and cultivated (Gertis,1925). Deforestation of the Palau Penida Archipelago made primary and secondary forest, vital avian habitats, fragmented and in short supply. As of 2017, the largest remaining forested portion of the island is less than ten hectares in area, centrally located on the island, at the top of Mount Mundi, an area with slopes up to a 40% grade that is transected by a single north-south road. These conditions exacerbate the fragmentation of forested areas and decrease the habitability of the island’s primary forests.

Intrinsic physical and biological challenges to avifaunal biodiversity on islands have historically been exacerbated by human presence (Brown 1997). Avian biodiversity can serve as a proxy for overall diversity (Lees et al. 2012, Moura et al. 2013). Birds are one of the largest vertebrates on these islands and serve as ecosystem engineers, seed dispersers, insectivores, and as pollinators (Whelan et al. 2008). Invasive species can imperil a productive avian community if their presence diminishes endemic biodiversity and affect the aforementioned ecosystem roles. The Indonesian Palau Penida Archipelago, host to several threatened and endangered species, is a release site for the critically endangered Bali Starling, *Leucopsar rothschildi*, despite never having a formal avifaunal biodiversity survey conducted on the islands (Juniartha, 2007). This preliminary survey can serve as a tool to inform future conservation management decisions in hopes of maintaining biodiversity on the ancillary reserve that is Indonesia’s Palau Penida Archipelago.

## MATERIALS & METHODS

### Study Area

This study was conducted on the islands of Nusa Penida and Nusa Lembongan, part of Bali’s Klungkung Regency. Nusa Penida, the largest of the three islands, is located at 8.727°S, 115.544°E. Nusa Lembongan is accessible by foot from Nusa Ceningan and 600 meters of water separates them from Nusa Penida. Nusa Penida’s overall elevation ranges from 0 to 529 meters, where Nusa Lembongan peaks at 29 meters.

### Bird Sampling

Thirty-two line transects were created throughout the islands based upon accessibility of the area and type of land use. I attempted to sample equal numbers of transects for each land use category, but areas defined as *Forest* for this study were sparse (Figure 1). Accessibility was determined by my ability to walk the line transect safely with little impeding the path, avoiding physical barriers that could interfere with wildlife observation. Transect elevation ranged from 5 meters in the mangroves of *Forest* transect 18 to 503 meters of the *Developed* transect 31 at the temple Puncak Mundi. Different land uses were found across the range of elevations on the islands. Each transect was surveyed over the month of May 2017, the second month of the dry season. Average temperature during the study was 27.5°C, which is in keeping with the recent historical mean during the same month. Precipitation occurred on three days during the study; 120 mm is the current average precipitation for the month of May (University of Maine 2017).

**Figure 1:**
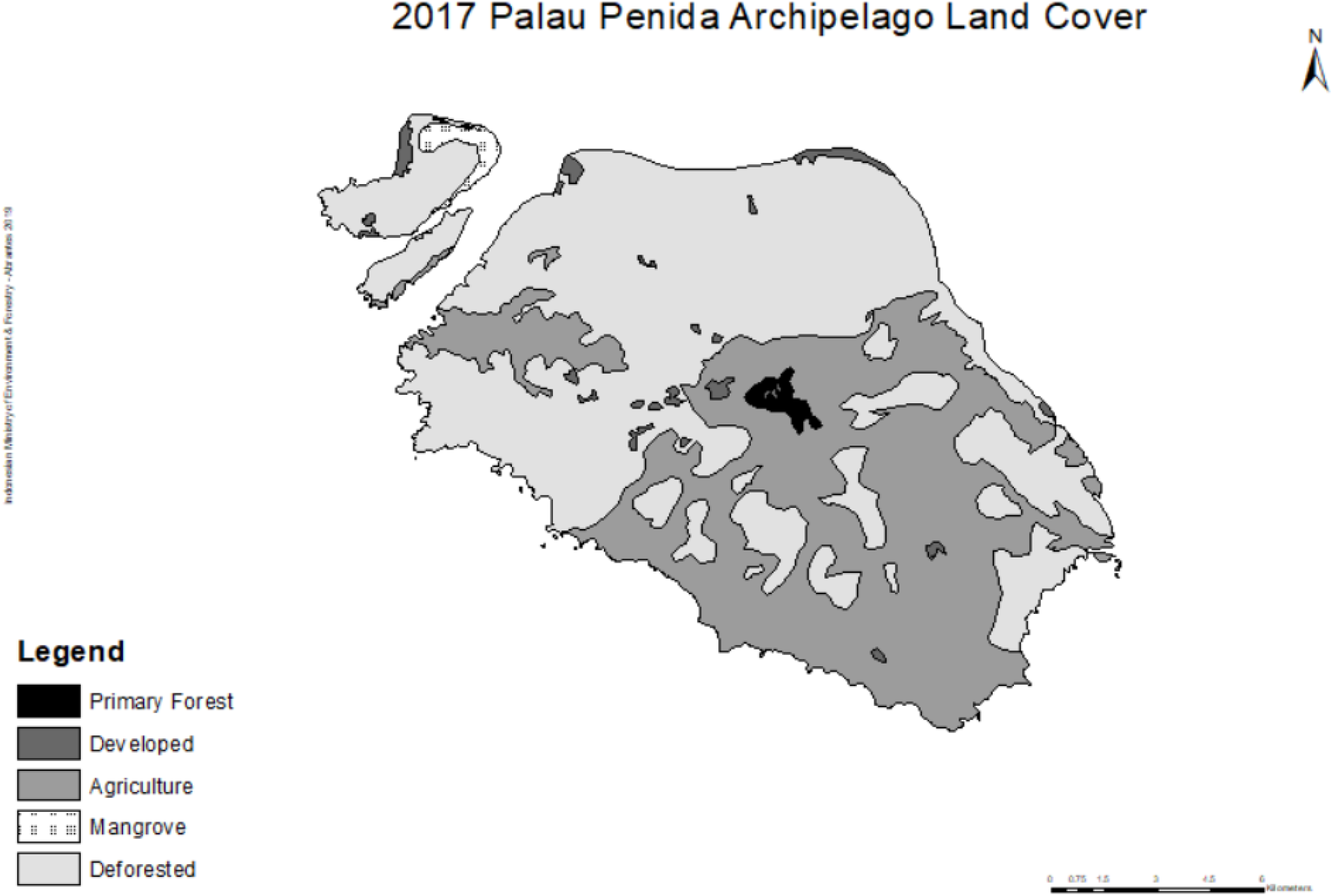
Map of Land Cover for Nusa Penida, Nusa Ceningan, and Nusa Lembongan, 2017. Remaining primary forest, in black, is also the highest elevation of the three islands.

Each unique 30-meter-wide transect was measured and marked with flagging tape so I could quickly identify the survey location upon return in the evening. Transects were walked down the middle, with 15 meters on each side, to a length of 250 meters as measured by GPS from a Garmin 735XT (Garmin Ltd., Lenexa, KS) for a total area of 7500 m^2^ per transect. Surveys focused on Nusa Penida, but two sites were chosen on Nusa Lembongan because a Bali Starling population was recently reported there by the Begawan Foundation (Halaouate, 2015). Transects were completed within 30 minutes, accounting for pauses to count, confirm identification, or scale terrain. The 32 transects were cumulatively surveyed 60 times; four transects were surveyed only once due to flooding of the only access routes (Table 1).

**Table 1(a).**
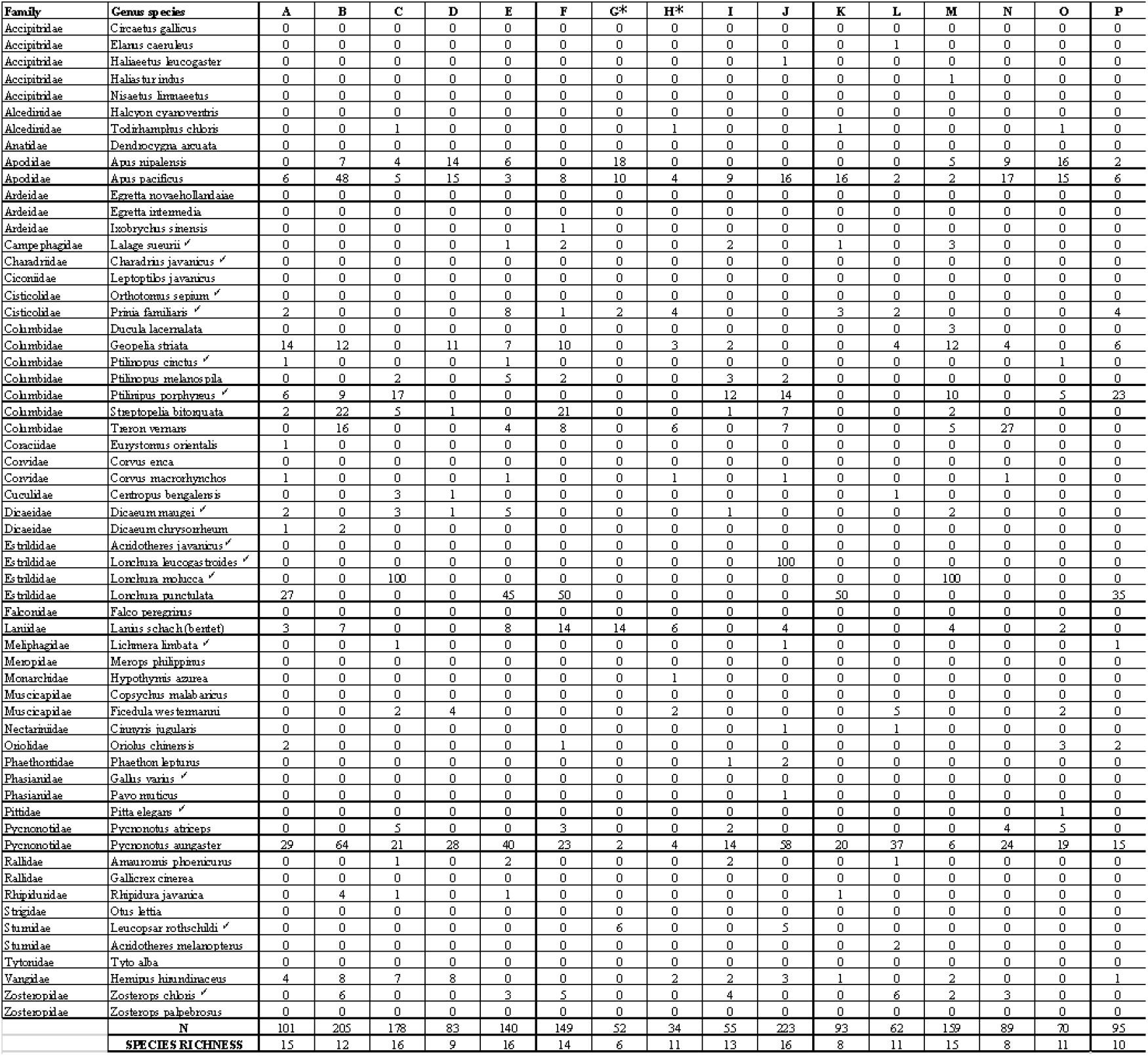
Bird counts from surveys (A-P) of the Palau Penida Archipelago, May 2017. Checkmarks (✓) indicate species listed as endemic peravibase.org. * Are survey transects that were not repeated due to weather.

**Table 1b:**
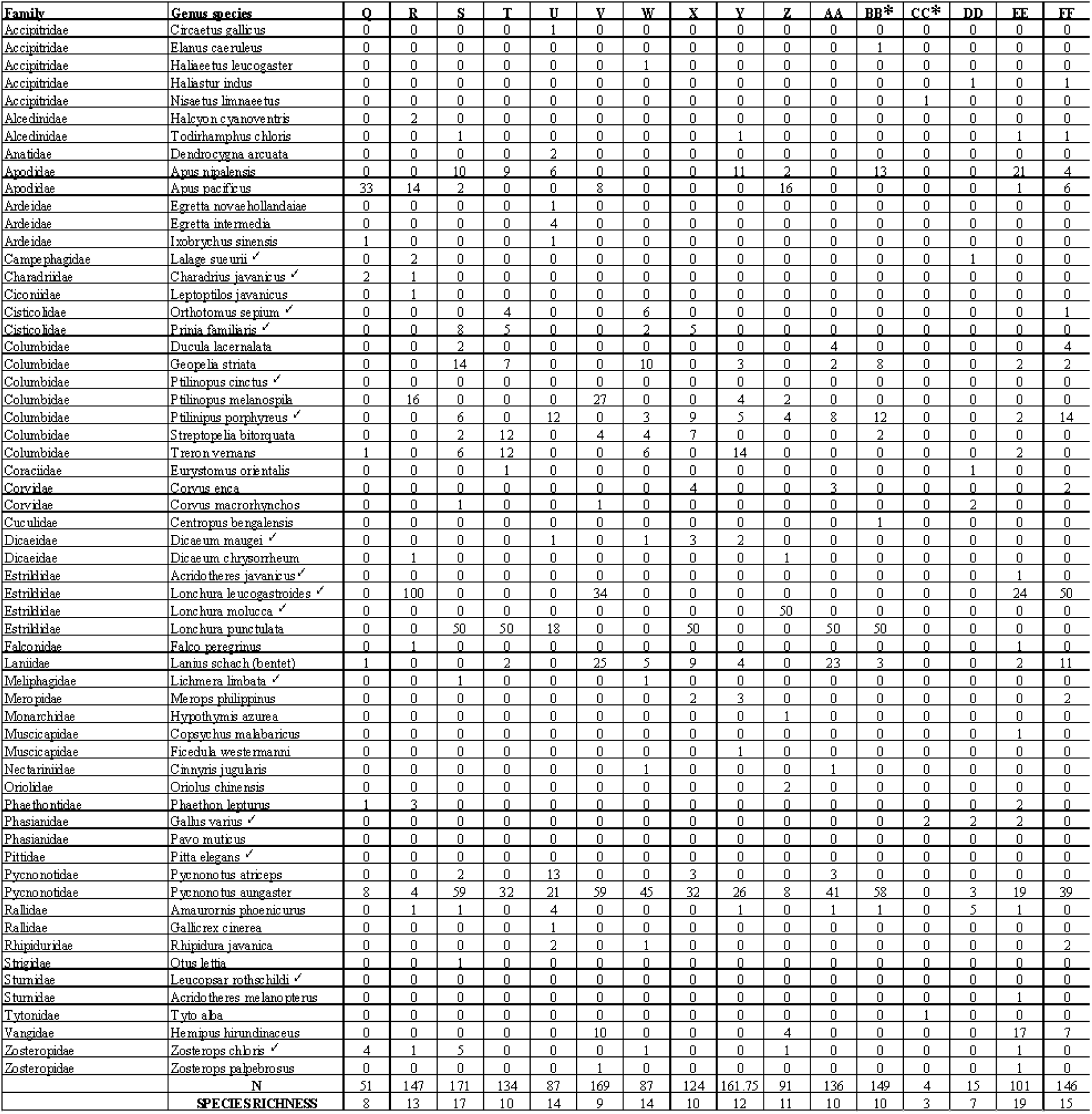
Bird counts from surveys O-FF of the Palau Penida Archipelago, May 2017. ✓ Native per avibase.org. *Site visited only once due to weather.

Surveys commenced between 0600-0730 and 1700-1830 to account for the matutinal and crepuscular nature of bird activity. I identified birds by sight only. All birds were counted individually in the field, except for flocks of *Lonchura*. These flocks were initially counted via video and numbers were estimated for flocks with greater than 50 members. Fly-over birds, those clearing the canopy or tallest tree in the transect, were not counted per protocol dictated in previous avifaunal studies (Lees et al. 2012, Moura et al. 2013). Identification was made on site by two observers, walking together, with the aid of 10x binoculars and a Nikon DSLR camera using an 18-55mm lens (Nikon, Tokyo, Japan).

### Land Use Designation

Land use categories *Agriculture, Deforested, Developed*, and *Forest* were appointed to every transect. Transects that are currently cultivated for crops such as cassava, banana, and corn were designated *Agriculture. Deforested* land was similar, but these transects were not as close to domiciles and showed no recent signs of disturbance beyond the established century old stone-terraced farming steps. Banana trees, coconut palms, various shrubs and grasses were regularly found among these transects. Land that has distinctly been developed for tourism, has recently been cleared for home expansion, or tracts of limestone excavation composed *Developed* transects. These had the sparsest flora, typically shrubs and grasses with trees on the periphery. Designation as *Forest* was limited to primary or secondary forest that had not been destroyed, replanted, or significantly modified over the last century. Mature mangrove forest found on Nusa Lembongan was also included in this category. Teak, Banyan, bamboo, palm trees, and very dense shrubs littered these limited landscapes. Land cover was not uniform among all transects of a given land use category. Quantity of transect type varied based upon perceived prevalence of that land use across the islands. Ten *Agriculture*, nine *Deforested*, eight *Developed*, and five *Forest* transects were evaluated during this study.

### Endemic Designation

Avibase (avibase.org) was used as a guide to define the endemic versus exotic species of Bali and its Klunkung Regency. Of the 410 bird species listed for Bali in the database, the Bali Starling or Bali Myna (*Leucopsar rothschildi*) is the sole endemic bird listed as unique to Bali itself. Because of the proximity to neighboring islands Java and Lombok and the study’s focus on a volant organism, the 38-species labeled “endemic (country/region)” in the database are considered endemic for the purposes of this study.

### Data Analysis

Endemic species richness, exotic species richness, density, and evenness were quantified for each survey. Simpson and Shannon-Weiner indices were used to calculate biodiversity. I calculated species density of individual birds for each meter squared per transect. Jaccard indices were used to quantify similarities in the bird communities among different land use categories. Jaccard similarity coefficients were calculated by comparing the presence of every species across all transects in two land use categories, shown as a percentage of species in common, divided by total unique species found in each land use comparison. Statistical examination of biodiversity indices and subcomponents across land use categories was done using a one-way analysis of variance (ANOVA) with a 95% confidence interval. Treatment effect for values of significance (p < 0.05) was followed by a Tukey post-hoc test. We created a Pearson Correlation Matrix to examine relationships between variables not analyzed via ANOVA. Statistical analysis was executed on SPSS Version 24 (IBM, New Castle, NY).

## RESULTS

### Species Richness

I counted 3,475 birds throughout the study, observed among 60 species across 32 transects on Nusa Penida and Nusa Lembongan. The most species rich surveys were dawn transects 1 (*Deforested*; 44 meters elevation), 5 (*Deforested;* 33 meters), 10 (*Developed;* 68 meters), and dusk transect 3 (*Developed*; 16 meters); all with 12 species each. The least species rich survey was transect 17 (*Forest;* 429 meters) at dusk with only two species present, the fork-tailed swift *(Apus pacificus)* and sooty-headed bulbul (*Pycnonotus aurigaster). Deforested* transects had the greatest mean overall species richness (Table 2; 7.71 ± 2.82) and *Forest* transects had the lowest (Table 2; 5.86 ± 3.31) but these trends were not significant (p=0.23). All individual survey data can be found in Table 1.

**Table 2:** Time of survey (AM/PM), Land use type (A=Agri culture, D=Developed, F=Forest, T=Deforested), Elevation in meters, Endemic species richness, and exotic species richness, by transect.

| Transect | Time | LandUse | Elevation | EN Rich | EX Rich |
| --- | --- | --- | --- | --- | --- |
| 1 | A | T | 44 | 3 | 9 |
| 2 | A | T | 121 | 0 | 9 |
| 1 | P | T | 44 | 0 | 7 |
| 2 | P | T | 121 | 1 | 10 |
| 3 | A | D | 16 | 2 | 7 |
| 4 | A | D | 83 | 1 | 5 |
| 3 | P | D | 16 | 3 | 9 |
| 4 | P | D | 83 | 0 | 6 |
| 5 | A | T | 33 | 5 | 7 |
| 6 | A | A | 69 | 2 | 3 |
| 5 | P | T | 33 | 2 | 7 |
| 6 | P | A | 69 | 1 | 10 |
| 7 | A | D | 170 | 2 | 4 |
| 8 | A | A | 386 | 1 | 10 |
| 9 | A | F | 91 | 2 | 6 |
| 10 | A | D | 68 | 2 | 10 |
| 9 | P | F | 91 | 1 | 5 |
| 10 | P | D | 68 | 2 | 7 |
| 11 | A | D | 50 | 1 | 4 |
| 12 | A | D | 374 | 2 | 5 |
| 11 | P | D | 50 | 1 | 4 |
| 12 | P | D | 374 | 0 | 6 |
| 13 | A | A | 154 | 3 | 8 |
| 14 | A | A | 14 | 0 | 5 |
| 13 | P | A | 154 | 2 | 5 |
| 14 | P | A | 14 | 1 | 5 |
| 15 | A | T | 276 | 1 | 3 |
| 16 | A | T | 231 | 0 | 4 |
| 15 | P | T | 276 | 1 | 8 |
| 16 | P | T | 231 | 2 | 6 |
| 17 | A | F | 429 | 2 | 6 |
| 18 | A | F | 5 | 4 | 6 |
| 17 | P | F | 429 | 0 | 2 |
| 18 | P | F | 5 | 1 | 4 |
| 19 | A | A | 242 | 1 | 7 |
| 20 | A | A | 14 | 2 | 8 |
| 19 | P | A | 242 | 2 | 8 |
| 20 | P | A | 14 | 1 | 3 |
| 21 | A | T | 244 | 1 | 5 |
| 22 | A | D | 107 | 0 | 8 |
| 21 | P | T | 244 | 0 | 10 |
| 22 | P | D | 107 | 1 | 4 |
| 23 | A | A | 51 | 3 | 5 |
| 24 | A | A | 228 | 2 | 5 |
| 23 | P | A | 51 | 2 | 6 |
| 24 | P | A | 228 | 0 | 5 |
| 25 | A | T | 123 | 0 | 6 |
| 26 | A | T | 132 | 1 | 6 |
| 25 | P | T | 123 | 1 | 6 |
| 26 | P | T | 132 | 1 | 5 |
| 27 | A | A | 178 | 0 | 9 |
| 28 | A | A | 282 | 0 | 6 |
| 27 | P | A | 178 | 0 | 4 |
| 28 | P | A | 282 | 0 | 5 |
| 29 | A | F | 192 | 1 | 2 |
| 30 | P | F | 488 | 2 | 5 |
| 31 | A | D | 503 | 1 | 9 |
| 31 | P | D | 503 | 3 | 7 |
| 32 | A | T | 20 | 1 | 6 |
| 32 | P | T | 20 | 1 | 8 |

Forty-three species were represented in the 1793 birds counted during morning surveys (mean overall richness = 7.82, ±2.42) versus the 1682 birds among 56 species (mean overall richness = 7.16, ±2.50) in evening transects. I counted fourteen species (23%) categorized as endemic to Bali and the Nusa Islands. Of the 3,475 birds counted, 719 (20.69%) were endemic. However, only two species, the Javan munia (*Lonchura leucogastroides)* and the black-faced munia (*Lonchura molucca)*, accounted for 77.61% (558 of 719) of the individuals of these endemic species. Among the exotic species, the invasive sooty-headed bulbul *(Pycnonotus aurigaster)* was nearly ubiquitous throughout the island, noted in 30 of 32 transects (94%). Sooty-headed bulbuls accounted for 31.13% (858 of 2756) of all individual invasive birds and 24.69% (858 of 3475) of the total individuals across all surveys. Eight transects (25%) did not have a single endemic bird present (Table 1).

Differences in exotic species richness among land use type was marginally significant (F3,28 = 2.91, p = 0.052). The strongest distinction in exotic richness was among *Forest* and *Developed* transects with *Forest* transects having four times lower exotic species richness, on average, comparatively. Excluding the two *Lonchura* species, the total of all 161 individual endemic birds account for only 4.63% of total birds surveyed. Endemic species richness was not significantly different across land use categories (F3,28 = 0.14, p = 0.94). However, endemic species richness and exotic species richness were significantly positively correlated (Table 2). Repeated measures ANOVA showed that species richness of exotics (9.2 ± 0.5) was consistently greater than that of endemic species (2.3 ± 0.2) (Table 3; F1,28 = 252.93, p < 0.0001). We did not find a significant relationship between overall species richness and land use (F3,28 = 1.38, p = 0.27)

**Table 3.** Survey Diversity: Shannon diversity index, Simpson diversity index, species density, and species evenness, by transect from May 2017 on the Palau Penida Archipelago.

| Transect | Time | Shannon | Simpson | Density | Evenness |
| --- | --- | --- | --- | --- | --- |
| 1 | A | 1.860986 | 0.758065 | 0.008267 | 0.748916 |
| 2 | A | 1.860092 | 0.801541 | 0.013067 | 0.846564 |
| 1 | P | 1.547532 | 0.717949 | 0.0052 | 0.795274 |
| 2 | P | 2.040337 | 0.282051 | 0.014267 | 0.850886 |
| 3 | A | 1.228167 | 0.565463 | 0.0108 | 0.558963 |
| 4 | A | 1.389715 | 0.706667 | 0.004 | 0.775615 |
| 3 | P | 1.673366 | 0.691253 | 0.012933 | 0.673412 |
| 4 | P | 1.6184 | 0.780349 | 0.007067 | 0.903246 |
| 5 | A | 1.632656 | 0.710948 | 0.0132 | 0.657029 |
| 6 | A | 0.986631 | 0.508089 | 0.01 | 0.613028 |
| 5 | P | 1.899234 | 0.811422 | 0.005467 | 0.864379 |
| 6 | P | 2.061339 | 0.844412 | 0.009867 | 0.859645 |
| 7 | A | 1.53735 | 0.754438 | 0.006933 | 0.858011 |
| 8 | A | 2.226211 | 0.878893 | 0.004533 | 0.928402 |
| 9 | A | 1.846997 | 0.805293 | 0.003067 | 0.888218 |
| 10 | A | 1.643155 | 0.729529 | 0.0168 | 0.661254 |
| 9 | P | 1.493328 | 0.736328 | 0.004267 | 0.833442 |
| 10 | P | 1.505936 | 0.677862 | 0.012933 | 0.685381 |
| 11 | A | 1.07787 | 0.571055 | 0.011067 | 0.669719 |
| 12 | A | 1.349593 | 0.621958 | 0.005733 | 0.693554 |
| 11 | P | 1.359237 | 0.68 | 0.001333 | 0.844541 |
| 12 | P | 1.229099 | 0.570637 | 0.002533 | 0.685973 |
| 13 | A | 1.473434 | 0.583881 | 0.010533 | 0.61447 |
| 14 | A | 1.237812 | 0.650888 | 0.0052 | 0.769096 |
| 13 | P | 1.241433 | 0.573438 | 0.010667 | 0.63797 |
| 14 | P | 1.627407 | 0.7792 | 0.006667 | 0.908273 |
| 15 | A | 1.18582 | 0.660494 | 0.0024 | 0.855389 |
| 16 | A | 0.953964 | 0.523148 | 0.0048 | 0.68814 |
| 15 | P | 1.847777 | 0.808432 | 0.006933 | 0.84096 |
| 16 | P | 1.392538 | 0.61649 | 0.007867 | 0.669669 |
| 17 | A | 1.211197 | 0.523009 | 0.004933 | 0.55124 |
| 18 | A | 1.29768 | 0.476991 | 0.010933 | 0.563575 |
| 17 | P | 0.682908 | 0.489796 | 0.001867 | 0.985228 |
| 18 | P | 0.788875 | 0.385799 | 0.008667 | 0.490156 |
| 19 | A | 1.484903 | 0.710139 | 0.016 | 0.714088 |
| 20 | A | 1.716853 | 0.727982 | 0.014 | 0.74562 |
| 19 | P | 1.724 | 0.747405 | 0.0068 | 0.748724 |
| 20 | P | 0.824539 | 0.561237 | 0.003867 | 0.594779 |
| 21 | A | 1.196711 | 0.569444 | 0.0032 | 0.667897 |
| 22 | A | 1.572247 | 0.73875 | 0.010667 | 0.756091 |
| 21 | P | 1.99078 | 0.833963 | 0.0084 | 0.864585 |
| 22 | P | 1.418785 | 0.724656 | 0.011867 | 0.881541 |
| 23 | A | 1.858639 | 0.820452 | 0.003867 | 0.893816 |
| 24 | A | 1.739204 | 0.786704 | 0.005067 | 0.893774 |
| 23 | P | 1.227928 | 0.559453 | 0.007733 | 0.590508 |
| 24 | P | 1.170504 | 0.600054 | 0.011467 | 0.727275 |
| 25 | A | 1.289389 | 0.662037 | 0.0048 | 0.719622 |
| 26 | A | 0.973947 | 0.41506 | 0.0088 | 0.50051 |
| 25 | P | 1.720225 | 0.78238 | 0.0052 | 0.884021 |
| 26 | P | 1.25553 | 0.6176 | 0.003333 | 0.700724 |
| 27 | A | 1.522386 | 0.693841 | 0.014133 | 0.692868 |
| 28 | A | 1.289418 | 0.640595 | 0.012267 | 0.719638 |
| 27 | P | 1.011949 | 0.588889 | 0.004 | 0.729967 |
| 28 | P | 0.845055 | 0.453678 | 0.0076 | 0.525062 |
| 29 | A | 1.039721 | 0.625 | 0.000533 | 0.946395 |
| 30 | P | 1.767009 | 0.8 | 0.002 | 0.908063 |
| 31 | A | 1.657339 | 0.685547 | 0.004267 | 0.719773 |
| 31 | P | 1.612193 | 0.746531 | 0.0092 | 0.700167 |
| 32 | A | 1.418306 | 0.676177 | 0.013067 | 0.728865 |
| 32 | P | 1.672012 | 0.736111 | 0.0064 | 0.760966 |

Land-use category had a significant effect on the difference between endemic and exotic species richness (F3,24 = 3.89, p < 0.02), again with *Forest* having substantially lower exotics than any other transect type. There was no significant effect of land use category, species density (F3,28 = 2.29, p = 0.10), or species evenness (F3,28 = 0.27, p = 0.85), though species density significantly decreased with elevation (Table 1). Overall, we found no significant difference in cumulative species richness between morning and evening transects (Table 2; F1,58 = 0.34, p = 0.57).

### Species Diversity

Among diversity indices, there was a significant relationship between land-use type and the Shannon-Weiner diversity index (Table 3; F3,28 = 3.18, p = 0.039). The most substantial difference, again, was between Shannon indices for *Deforested* and *Forest* transects with *Forest* transects being significantly less biodiverse (Tukey p < 0.05). However, there was no significant relationship between land use category and the Simpson index (Table 3; F3,28 = 0.97, P = 0.43). Shannon diversity was positively correlated (p < 0.05) with both overall species richness and evenness (Table 2) whereas Simpson diversity was correlated with evenness but not overall richness (Table 2). *Deforested* transects had the greatest mean Shannon diversity (Table 3; 1.52 ± 0.49) and the lowest mean Simpson diversity (Table 1; 0.58 ± 0.25). We found the lowest mean Shannon diversity among *Forest* transects (Table 3; 1.18 ± 0.55) where the highest Simpson diversity mean was in *Developed* transects (Table 3; 0.691 ± 0.19). There was no significant difference between transects at different times of day for the Shannon index (Table 3; F1,58 = 0.012, P=0.91), Simpson index (Table 3; F1,58=0.20, P=0.66), or evenness (Table 3; F1,58 = 0.53, P = 0.47).

### Jaccard Coefficient of Similarity

*Deforested* and *Agriculture* transects had the most similar avian communities, with a Jaccard coefficient of 68.09 (Table 4), and a greater proportional similarity despite having the fewest total species in common, 47. Forest and Agriculture had the least similarity with the lowest Jaccard index, 38.30 (Table 4), and 48 species in common.

## DISCUSSION

By count alone, exotic birds observed throughout this study dramatically outnumbered endemic species nearly fourfold (2756:719), however proportion does not paint a complete picture of the avifauna of these Indonesian islands. Two species, *Lonchura leucogastroides* and *Lonchura molucca*, accounted for greater than three quarters of all observed birds categorized as endemic. Examining total species, only 60 (14.63%) of the 410 of the previously documented endemic and exotic species of birds were observed.

Most striking among the results was the low species density in *Forest* transects. Intuitively, forests make a likely candidate for avian habitat, including for the Bali Starling. Forested areas at higher elevations remain unlikely sites for development and agriculture due to the steep slope and heavy wind exposure, the same reasons most endemic and exotic birds alike are not making their homes above 250 meters (Table 2). Of the 11 individual critically endangered Bali starlings observed, none were found in *Forest* transects.

Comparison of surveys at dawn and dusk showed more matutinal activity across surveys by more individual birds, though fewer species. Mornings on the Palau Penida Archipelago are cooler and moister. Birds are still present and active at dusk, though without midday precipitation, morning conditions are likely more ideal for bird activity than the hot dry evenings after hours of near-Equatorial sun.

Higher overall species richness was observed at transects of lower elevation, with the greatest species richness at transects below 50 meters. *Forest* fragments account for the highest and lowest points on these islands. Deforestation accompanied by changes in land use and mixed vegetative cover have opened the door for invasive species, particularly those with adaptations to thrive in Nusa Penida’s locally-reported worsening dry season. This exacerbates the challenges imposed by the Palau Penida Archipelago’s natural hydrogeology (Giambelli 1999).

Exotic species such as the frugivorous sooty-headed bulbul are outcompeting the rarer endemic birds. Papaya, a favorite bulbul food native to the Americas, are found throughout the islands benefiting invasive birds who thrive on them. Dominance by exotic bird species is also likely to indicate the prevalence of exotic species in other genera, such as the seeds of non-native vegetation invasive birds can and have spread (Dawson et al. 2017). Reforestation, when well-monitored and protected, stands to promote endemic bird survival and overall island biodiversity. Small reforestation attempts have been made and continue to be made on Nusa Penida though monitoring the trees post-sapling is inconsistent.

The Bali Starling was the only IUCN red-list endangered or critically endangered bird observed throughout this study, despite the six other critically endangered species of bird previously documented on Nusa Penida (avibase.org). Observation of 11 Bali starlings is a beacon of hope, though meager compared to the 2015 Begawan survey reporting several times the Bali Starling population (Halaouate 2015). Unfortunately, these and other birds face threats beyond habitat destruction. Those living around agricultural or developed areas can easily be captured and exported by boat from Bali and the Nusa’s to Lombok, Java, and beyond to be sold with the steep price tag their rarity brings (Jepsen 2016). As exotic species continue to adapt to the conditions on the Palau Penida Archipelago, outcompeting and exploiting each island’s resources, the fate of many endemic species could potentially be imperiled.

Maintenance of biodiversity on the Palau Penida Archipelago is essential to the islands’ ecosystem and the many services it provides. Recently, the ancillary avian reserve was established on these islands in an attempt to promote biodiversity, preserve endangered species, and mitigate the significant, detrimental anthropogenic changes in land use of the recent past. Observations and correlations derived from this study are a gauge of the direction of land use reflecting on biodiversity of the landscape, not an absolute count of populations of avian species, particularly the critically endangered Bali Starling. Further studies of the avian populations on these islands are likely to provide a model for human interference and the fate of Indonesia’s biodiversity.

---

**Figure 1** Map of 2017 land cover on the Palau Penida Archipelago.

**Table 1 (a) Bird counts** from surveys (A-P) of the Palau Penida Archipelago, May 2017. **(b)** Bird counts from surveys (Q-FF) of the Palau Penida Archipelago, May 2017. Checkmarks (✓) indicate species listed as endemic per avibase.org. *Are survey transects that were not repeated due to weather.

**Table 2 Survey** Time of survey (AM/PM), Land use type (A=Agriculture, D=Developed, F=Forest, T=Deforested), Elevation in meters, Endemic species richness, and exotic species richness, by transect, May 2017.

**Table 3 Survey Diversity** Shannon diversity index, Simpson diversity index, species density, and species evenness, by transect from May 2017 on the Palau Penida Archipelago

**Table 4 Jaccard Index** coefficients and species in common among land use categories for birds observed on the Palau Penida Archipelago, May 2017.

